# A Libra in the Brain: Neural Correlates of Error Awareness Predict Moral Wrongness and Guilt Proneness

**DOI:** 10.1101/2024.04.17.589679

**Authors:** Chenyi Chen, Scott S.-C. Kao, Yang-Teng Fan, Yu-Chun Chen, Yawei Cheng

**Affiliations:** Graduate Institute of Injury Prevention and Control, College of Public Health, Taipei Medical University, Taipei, Taiwan; Graduate Institute of Mind, Brain and Consciousness, College of Humanities and Social Sciences, Taipei, Taiwan; Department of Physical Medicine and Rehabilitation, Wan Fang Hospital, Taipei Medical University, Taipei, Taiwan; Psychiatric Research Center, Wan Fang Hospital, Taipei Medical University, Taipei, Taiwan; The Innovative and Translational Research Center for Brain Consciousness, Taipei Medical University, Taipei, Taiwan; Neuroscience Research Center, Taipei Medical University Hospital, Taipei, Taiwan; Institute of Neuroscience, National Yang Ming Chiao Tung University, Taipei, Taiwan; Graduate Institute of Medicine, Yuan Ze University, Taoyuan, Taiwan; Department of Physical Education, National Taiwan University of Sport, Taichung, Taiwan; Department of Physical Medicine and Rehabilitation, National Yang Ming Chiao Tung University Hospital, Yilan, Taiwan

**Author notes:** Corresponding Authors: Prof. Yawei Cheng Institute of Neuroscience National Yang Ming Chiao Tung University No. 155, Sec. 2, St. Linong, Dist. Beitou Taipei 112, Taiwan, ROC.

## Abstract

Error awareness is a fundamental mechanism in humans. Through traditional psychological tasks, certain neural activities that represent errors in objective manners have been identified. However, there is limited knowledge on how humans subjectively determine right from wrong in moral contexts. In this study, participants (N=39) mentally simulated themselves as the agents of moral and immoral behaviors, while viewing a series of actions with EEG recording and MRI scanning, respectively. A significant difference in error-related negativity (ERN) was observed among morally wrong scenarios, accompanied by higher wrongfulness ratings. Additionally, individual differences in guilt-proneness could predict the subjects’ ERN amplitude. The ERN amplitude was correlated with the BOLD activity in the anterior mid-cingulate cortex and anterior insula to immoral scenarios, reflecting error awareness toward moral wrongfulness. The late potential component displayed greater negativity to immoral scenarios and was correlated with BOLD activities in the amygdala, ventromedial prefrontal cortex, and temporoparietal junction, indicating cognitive and affective evaluation in moral judgment. In line with the moral dynamic framework, our results demonstrated individual variability in moral judgments, as indicated by dispersed and overlapping cognitive neural networks. This suggests that subjective evaluations of wrongfulness are underpinned by neural mechanisms, associated with those involved in objective error awareness.

## Introduction

Imagine witnessing a flash mob robbery on the streets, ransacking every store. This startling situation is not only alarming but also clearly recognized as illegal and wrong. The neural mechanisms through which humans detect and process such errors have been the subject of study for decades in cognitive neuroscience.

Error-related negativity (ERN), a critical indicator of the early processing of errors, has been extensively examined. Since Falkenstein and colleagues (1991) first identified frontocentral negativity in event-related potentials (ERPs) following error trials in 1991, a wealth of research has confirmed its relationship with conflict monitoring (Kirschner, Humann, Derrfuss, Danielmeier, & Ullsperger, 2021; Markus Ullsperger, Fischer, Nigbur, & Endrass, 2014). It is argued that ERN involves the conscious awareness of errors, as demonstrated through traditional psychological paradigms like the Go/No-go and flanker tasks (Imburgio et al., 2020; Kirschner et al., 2021; M. Scheffers & Coles, 2000; M. K. Scheffers, Coles, Bernstein, Gehring, & Donchin, 1996; Wang, Gu, Zhao, & Chen, 2020; Wessel, 2012). Interestingly, ERN is also elicited in error trials using affective stimuli (Suzuki, Ait Oumeziane, Novak, Samuel, & Foti, 2020). Typically peaking around 50-150 milliseconds post-error (Falkenstein et al., 1991; Markus Ullsperger, Danielmeier, & Jocham, 2014), ERN amplitudes are prominently larger in frontal and central electrodes(M. Scheffers & Coles, 2000). The anterior cingulate cortex is presumed to generate ERN, with supporting evidence from LORETA source localization studies (Herrmann, Römmler, Ehlis, Heidrich, & Fallgatter, 2004) and the direct coupling of fMRI/EEG results (Debener et al., 2005).

In real-life situations, unlike controlled experimental settings, we encounter complex circumstances that challenge our moral judgments. These judgments are shaped by ethical frameworks, such as utilitarianism or deontology, influencing how we approach dilemmas like the trolley problem (Greene et al., 2009; Malle, 2021). Contrary to the straightforward categorizations of right or wrong often seen in error-related experiments (M. K. Scheffers et al., 1996), the nuances of real-world moral decisions rarely fit such binary distinctions. Studies indicate that morality operates more as a dynamic system, influenced by a variety of factors (Moll, Zahn, de Oliveira-Souza, Krueger, & Grafman, 2005; Van Bavel, FeldmanHall, & Mende-Siedlecki, 2015). These include the severity of moral transgressions (Huang, Stanley, & De Brigard, 2020) and individual differences in moral judgment (Earp, McLoughlin, Monrad, Clark, & Crockett, 2021; Jin & Peng, 2021; Prehn et al., 2008). Intentional harm, widely recognized as morally reprehensible (K. J. Yoder & J. J. J. o. N. Decety, 2014), underscores the complexity of moral processing, which engages various brain regions such as the ventromedial prefrontal cortex (vmPFC), orbitofrontal cortex (OFC), dorsolateral prefrontal cortex (dlPFC), anterior cingulate cortex (ACC), temporoparietal junction (TPJ), and superior temporal sulcus (STS) (C. Chen, Martinez, Chen, Fan, & Cheng, 2022; Chenyi Chen, Martínez, & Cheng, 2020; de Oliveira-Souza, Zahn, & Moll, 2015).

On the other hand, guilt is a common moral emotion experienced in response to wrongdoing. This emotion is closely tied to moral judgments and can sometimes lead to prosocial behavior, encouraging actions that benefit others (de Hooge, Zeelenberg, & Breugelmans, 2007; Migliore et al., 2019; Tangney, Stuewig, & Mashek, 2007; Yang, Guo, Wu, & Robinson, 2022). Guilt-proneness, a personality trait predisposing individuals to feel negative emotions after committing wrongdoing (Cohen, Panter, & Turan, 2012), has been linked to a decreased frequency of immoral actions, such as lying (Cohen, Wolf, Panter, & Insko, 2011) The brain processes guilt in areas that partially overlap with those involved in the moral reasoning of harm. These areas include the medial prefrontal gyrus (MFG), anterior cingulate cortex (ACC), temporoparietal junction (TPJ), amygdala, and anterior insula (aIN), which integrate to give rise to the sensation of guilt (Cheng, Chou, Martinez, Fan, & Chen, 2021; Gifuni, Kendal, & Jollant, 2017). The distinct neural processes of guilt can also be explored through event-related potentials (ERPs), with late components reflecting cognitive processes such as appraisal of the situation or processing of emotions (MacNamara, Foti, & Hajcak, 2009; K. J. Yoder & J. Decety, 2014). Specifically, a late potential (LP) component observed in the frontal area during scenarios where individuals recognize their wrongdoing has been documented (Leng, Wang, Cao, & Li, 2017; Sanchez-Garcia et al., 2019).

Notably, while extensive research has shed light on how humans process errors using absolute and objective criteria of correctness, to date, no study has examined how the human brain integrates error awareness into the evaluation of subjective moral judgments, leaving the mechanisms involved unclear. Building upon our moral-action mental simulation task (C. Chen et al., 2022; Cheng et al., 2021), we have developed a paradigm involving animations that require participants to envision themselves as agents in scenarios that may or may not involve causing harm to others, accompanied by a subjective moral evaluation of the actions’ wrongness. Here, the study was to determine whether the ERP component known as Error-Related Negativity (ERN), typically associated with objective wrongness, could similarly reflect subjective moral judgments, mirroring its role in objective error awareness, and whether it could predict individual differences in guilt proneness. Furthermore, we utilized functional MRI to measure whole-brain activity, aiming to explore the potential of ERN and Late Positives (LP) in predicting patterns of brain activity. Participants engaged in similar tasks in both EEG and fMRI settings, with minor adjustments made to accommodate the scanner’s requirements. Region of Interest (ROI) analysis, focusing on guilt and moral processing, examined the relationship between specific brain regions and error awareness (C. Chen et al., 2022; Chenyi Chen et al., 2020; Cheng et al., 2021). We hypothesized that participants would show increased ERN and LP amplitude in scenarios involving harm, accompanied by a heightened sense of guilt. Additionally, we expect that guilt-prone individuals would display increased ERN activity, and that both ERN and LP would correlate with increased activation in regions associated with guilt and moral evaluation during harming scenarios, such as the mid-cingulate cortex (aMCC), amygdala, orbitofrontal cortex (OFC), and the right temporoparietal junction (rTPJ).

## Methods

### 1. Participants

Thirty-nine (19 females) healthy subjects, aged between 20 and 30 years old (mean ± SD: 22.38 ± 2.09), were recruited to participate in the EEG and fMRI version of our moral-action mental simulation paradigm. A total of 10 subjects were excluded due to either unable to complete two studies (EEG and fMRI) in different days, lousy signal acquisition, distraction during a task, or having seen the face stimulus ahead of both experiments. In the end, 29 subjects (14 females) remained. All participants provided written informed consent and received monetary compensation. The study was approved by the Institutional Review Board of National Yang Ming Chiao Tung University (YM107123F).

### 2. Stimuli

Fifty-eight volunteer models, aged between 20 and 40 years old (29 females, mean ± SD: 25.95 ± 5.83), were recruited. All provided written informed consent for the legal use of their portraits in the experiments. Glasses, earrings, and other accessories were removed for the photo session, and none of the volunteers had mustaches or beards. The photos were taken against a black background, with volunteers instructed to maintain a neutral facial expression in all scenarios. Four sets of photos were captured, each depicting a different action involving either a hand (holding a fist or an open palm) or a tool (a needle or a cotton swab). The needle and fist scenarios depicted harming actions, while the cotton swab and palm scenarios depicted non-harming actions. Each set consisted of three photos: one showing only the face, another with the action near the face, and a third with the action touching the cheek. These photos were arranged to simulate an animation. All photos were resized to ensure uniformity, positioning the model’s eyes at the center of the screen when displayed. The photos were utilized in both fMRI and EEG experiments.

### 3. Procedures

All participants needed to complete the guilt and shame proneness (GASP) scale beforehand. Then, they underwent similar moral-action mental simulation task in EEG and fMRI scanners, respectively, in different days apart from two-week wash-out interval. The sequence was counterbalanced, with half of the participants doing the task in EEG first and the rest conducting the task in fMRI scanner. There was at least a two-week interval between the two experiments. Both experiments required participants to watch the animations from a first-person perspective and simulate themselves as the agent of the action. Later, participants judged their perceived action as good or bad, then rate their warm glow or guilt feelings on a 7-point visual analog scale.

### 4. Guilt and shame proneness (GASP) scale

The Guilt and Shame Proneness (GASP) scale, as outlined by (Cohen et al., 2011), assesses an individual’s tendency to experience guilt and shame across a variety of scenarios. This scale is divided into four subscales: two dedicated to measuring guilt and two for measuring shame. Within these, one subscale in each pair focuses on negative evaluations—specifically, guilt-related negative behavior evaluations (NBE) and shame-related negative self-evaluations (NSE)—while the other subscale targets the propensity for action tendencies, including guilt-driven reparative actions and shame-induced withdrawal actions. Participants are presented with scenarios in which they are the protagonists who have committed a transgression. They are then asked to rate, on a 7-point Likert scale (where 1 is ’very unlikely’ and 7 is ’very likely’), their likelihood of experiencing specific reactions. This methodological approach aims to quantify the complex emotions of guilt and shame in a structured and nuanced manner.

### 5. EEG task, apparatus, recording, and analysis

The moral-action mental simulation task was administered using E-Prime software (Psychology Software Tools, Inc., Pittsburgh, PA). Each trial began with a fixation display lasting for one second, followed by the presentation of the first picture. Participants were instructed to press a button to progress the animation. Unlike our previous studies that employed similar stimuli and designs, where participants were explicitly primed with instructions to mentally simulate the outcome of the forthcoming harming or helping action before each trial, in this study, participants were not informed about the nature of the action—harmful or helpful—they would be simulating at the start of each trial. This approach aimed to more effectively trigger error-related or wrongness-related awareness after the outcomes of these actions were revealed in the subsequent animations (C. Chen et al., 2022; Chenyi Chen et al., 2020; Cheng et al., 2021). They were asked to envision themselves in the given action from a first-person perspective. For this, they viewed the initial image of a morally charged mini clip and were then able to press a handheld button at their own pace, with no set duration limit, initiating the subsequent sequence of images that depicted the actions to their full conclusion with varying moral outcomes (either harmful or neutral). A digital signal was captured upon the display of the second image to mark the beginning of the ERP waveform analysis. The duration of the second image was 50 milliseconds, and the third image lasted 950 milliseconds. Following each trial, participants engaged with a visual analog scale at their own pace (see Fig. 1A). The experiment comprised 180 trials in total, evenly divided into two blocks, with all four conditions presented in a randomized sequence (harm: needle and fist vs. no-harm: Q-tip and palm), ensuring an equal number of trials for each condition.

**Fig. 1:**
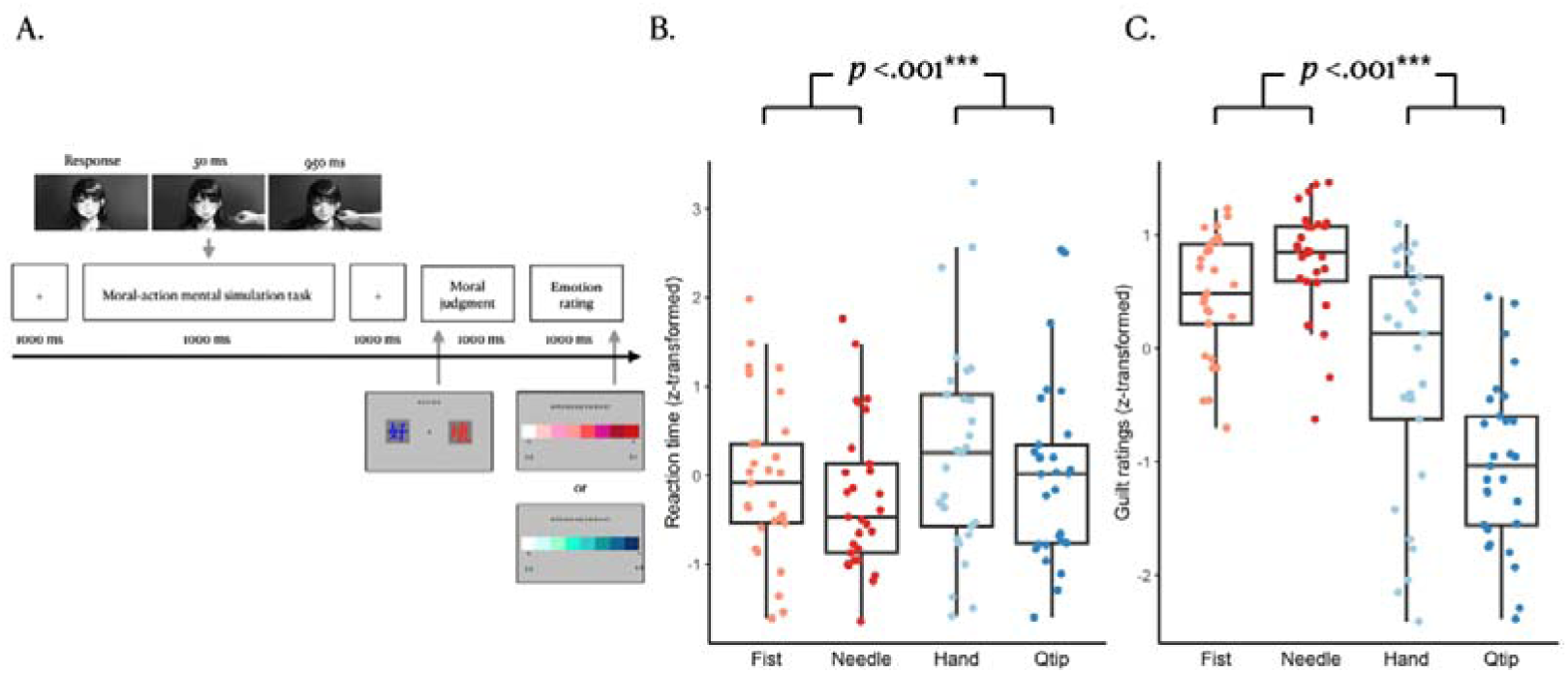
Experiment Design and Behavioral Results. **A.** A moral-action mental simulation task was designed, requiring participants to envision themselves performing the actions. **B.** A repeated-measures ANOVA was conducted to investigate the impact of harmfulness on participants’ responses. Both moral judgment reaction times and guilt ratings were normalized into z-scores. Participants assessed harming actions more quickly and reported a stronger sense of guilt (both p < .001).

Electrophysiological activity was recorded using a 32-channel cap following the International 10-20 system (EASYCAP, antiChamp, Brain Product). The online reference electrodes were positioned between Fz and Cz, with a ground electrode placed on the forehead. Additional electrodes were positioned below the left eye to monitor blinks and vertical eye movements. Electrode impedances were kept below 10 kΩ, and the online sampling rate was set at 1000 Hz.

Subsequently, all data were down-sampled to 512 Hz and filtered between 0.1 and 30 Hz for ERP analysis using the EEGLAB toolbox (v14.1.2) on MATLAB (MathWorks Inc., Sherborn, MA, USA). The data were re-referenced to the average of T9 and T10 and segmented into 500-millisecond epochs starting 200 milliseconds before the onset of the first picture. The baseline for correction was established using the average voltage 200 milliseconds prior to the response. Eye movements were corrected using Independent Component Analysis (ICA). Epochs exceeding ±75 μV, as well as sample-to-sample voltage changes exceeding 200 μV within 200 milliseconds, were excluded from analysis.

Electrophysiological activity for each condition was averaged across trials to generate event-related potentials (ERPs). The amplitude difference between 100 and 200 milliseconds post-stimulus was identified as the Error-Related Negativity (ERN) amplitude, and the activity between 300 and 400 milliseconds was designated as the Late Positive Potential (LPP) component. Data analysis focused on recordings from Fz and Cz. A three-way repeated measures ANOVA was conducted to examine the effects of harmfulness (harming vs. no-harming), utility (tool vs. hand), and recording site (Fz vs. Cz). Additionally, correlations were performed between individuals’ guilt ratings and their proneness to guilt.

### 6. fMRI task, scanning, preprocessing, and analysis

Stimuli were presented using E-prime software (Psychology Software Tools, Inc., Pittsburgh, PA) in conjunction with MRI-compatible goggles (VisualStim Controller, Resonance Technology Inc.). The moral-action mental simulation task was adapted for fMRI conditions. Initially, participants completed 60 trials inside the scanner, with 15 trials dedicated to each of the four conditions. The sequence of trials was both randomized and counterbalanced among participants. Upon the presentation of the first photo, participants were instructed to envision themselves as the acting agent and press a button to proceed with the animation. The durations for displaying the second and third photos were set at 500 milliseconds and 1500 milliseconds, respectively. A variable inter-trial interval of 2, 4, 6, 8, or 10 seconds was introduced to enhance data variability. Following the scanning session, ten images per condition, previously shown during scanning, were randomly selected for participants to assess their feelings of warmth or guilt, with this process also being randomized.

Imaging was conducted using a 3T Siemens Magnetom Trio-Tim system. High-resolution structural T1-weighted images were captured using a 3D magnetization-prepared rapid gradient echo (MP-RAGE) sequence (TR = 2530 ms, TE = 3.5 ms, FOV = 256 mm, flip angle = 7°, slice thickness = 1 mm, matrix = 256 × 256, no inter-slice gap). Functional images sensitive to changes in the blood oxygenation level-dependent (BOLD) T2* signal were acquired using a gradient echo-planar imaging (EPI) sequence (TR = 2200 ms, TE = 30 ms, FOV = 220 mm, flip angle = 90°, matrix = 64 × 64, voxel size = 3.4 × 3.4 × 3.0 mm, no inter-slice gap), obtaining 36 interleaved slices aligned with the anterior-posterior commissure line.

Preprocessing was executed using Statistical Parametric Mapping software (SPM12; Wellcome Trust Centre for Neuroimaging, UK) within MATLAB (MathWorks Inc., Sherborn, MA, USA). We coregistered structural T1 images with the mean functional images and generated a skull-stripped image from segmented gray matter, white matter, and cerebrospinal fluid (CSF) images. These segmented components were merged to form a subject-specific brain template. EPI images underwent realignment and were filtered using a 128-second cutoff before being coregistered to the brain templates and normalized to the Montreal Neurological Institute (MNI) space. Spatial smoothing was applied using an 8 mm full-width at half maximum (FWHM) isotropic Gaussian kernel.

For our analysis, we employed a general linear model (GLM) with two factors: harmfulness (harming vs. no-harming) and utility (tool vs. hand). A full factorial design was implemented for the analysis. At the second level, models were developed to explore brain activity correlated with guilt ratings, guilt-proneness, and event-related potential (ERP) activities (ERN and LPP) to elucidate individual differences in neural responses. Whole-brain exploratory analyses were adjusted for multiple comparisons using the family-wise error (FWE) rate, set at P < .05. Region of interest (ROI) analyses focused on eight predefined ROIs implicated in the processing of morality and guilt. Mean signal intensities (Beta weights) were extracted via MarsBaR (Brett, Anton, Valabrgue, & Poline, 2002) within SPM12. The selected ROIs were the amygdala, anterior insula (aIN), medial frontal gyrus (MFG), right posterior superior temporal sulcus/temporal parietal junction (pSTS/TPJ), orbitofrontal cortex (OFC), ventromedial prefrontal cortex (vmPFC), dorsolateral prefrontal cortex (dlPFC), and dorsal medial prefrontal cortex (dmPFC), referenced in studies by C. Chen et al. (C. Chen et al., 2022; Chenyi Chen et al., 2020; Cheng et al., 2021).

## Results

### 1. Moral judgments and subjective ratings of guilt

We conducted a two-way (action * utility) repeated-measures ANOVA to determine whether subjects responded and felt differently across various conditions. A significant interaction between harmfulness and utility was observed in moral judgments (F(1,28) = 5.76, p = .023, pη^2^= 0.171) and guilt ratings across conditions (F(1,28) = 35.55, p < .001, pη^2^= 0.559). In terms of moral judgments, participants assessed the harming actions as good or bad faster than the non-harming actions (F(1,28) = 17.84, p < .001, pη^2^= 0.389) (Fig. 1B). No significant difference was found in the type of utility (F(1,28) = 0.13, p = .727, pη^2^=0.004). Moreover, participants reported a significantly stronger sense of guilt when exposed to harming trials compared to non-harming trials (F(1,28) = 81.06, p < .001, pη^2^=0.743) (Fig. 1C). Tool scenarios also elicited a stronger sense of guilt compared to hand scenarios (F(1,28) = 16.49, p < .05, pη^2^= 0.149).

### 2. Electrophysiological results

Across the two electrodes (Fz and Cz), two negative waveforms were observed between 100 and 200 milliseconds and between 300 and 400 milliseconds (Fig. 2A). Data were collected using a fixed time window for all participants, setting the error-related negativity (ERN) amplitude between 110 and 150 milliseconds post-event and the late potential (LP) between 320 and 380 milliseconds. A three-way (harmfulness * utility * node) repeated-measures ANOVA was performed on the mean amplitudes of ERN and LP. For the ERN mean amplitude, significant differences were found in terms of harmfulness (F(1,28) = 4.90, p = .035, pη^2^= 0.149) and node (F(1,28) = 12.87, p = .001, pη^2^=0.315), with subjects showing a more negative response for harming than non-harming trials (Fig. 2A). No significant differences were observed for utility, nor were there any interaction effects. For the LP, significant differences were noted for harmfulness (F(1,28) = 44.20, p < .001, pη^2^= 0.612) and utility (F(1,28) = 33.52, p < .001, pη^2^= 0.545), with the mean amplitude for harming trials eliciting a larger response than non-harming trials across both electrodes. Tool scenarios also resulted in larger responses compared to hand scenarios. No significant differences were observed for nodes, and no interaction effects were found.

**Fig. 2:**
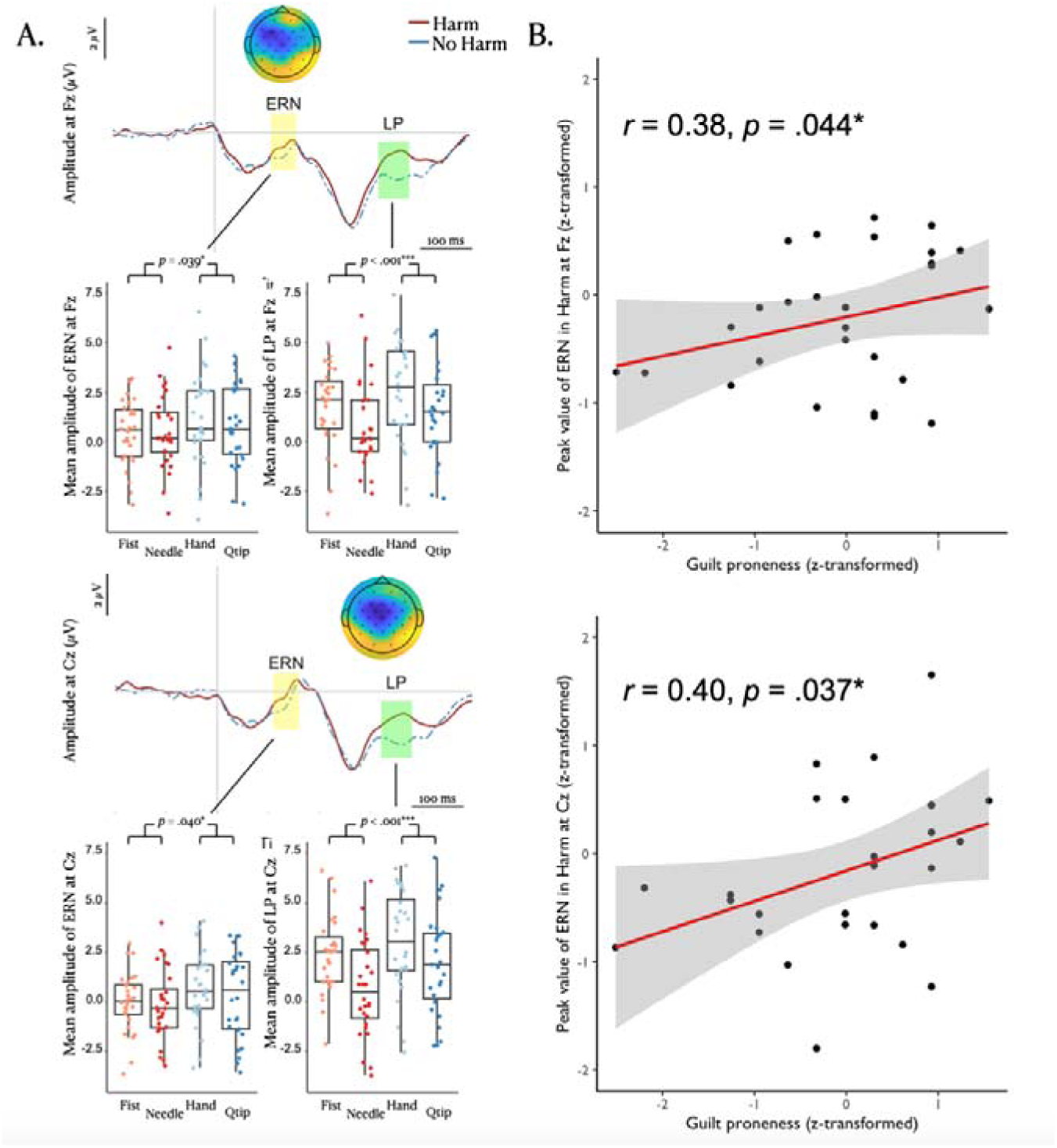
ERP results and the influence of individual guilt-proneness on ERN amplitude in harming trials. **A.** ERN and LP are established within fixed time windows for all participants. The top image illustrates amplitude measurements at Fz, while the bottom image does so at Cz. Subjects exhibited significantly more negative responses in both ERN (Fz: p = .039, Cz: p = .040) and LP (Fz: p < .001, Cz: p < .001) during harming trials. **B.** Participants’ guilt-proneness was found to significantly correlate with their peak ERN amplitude at Fz (r = .38, p = .044) and Cz (r = .40, p = .037). Individuals with higher levels of guilt-proneness displayed larger ERN amplitudes.

Correlation analysis was conducted to explore whether guilt-proneness or subjective guilt ratings influence participants’ electrophysiological responses in moral scenarios. We found that individuals with higher levels of guilt proneness exhibited larger ERN amplitudes. Specifically, the peak amplitude of ERN at Fz (r = 0.38, p = .044) and Cz (r = 0.40, p = .037) both significantly correlated with the participants’ guilt-proneness scores (Fig. 2B), but only in harming trials. The peak LP amplitude did not show any significant correlation with participants’ guilt proneness (all p > .05). Furthermore, neither ERN nor LP amplitudes significantly correlated with participants’ subjective guilt ratings (all p > .05).

### 3. fMRI results

Whole-brain analyses revealed no significant differences in neural activity, with a family-wise error (FWE) rate set at p < .05, irrespective of harmfulness or utility. Region of Interest (ROI) analyses, however, showed that compared to non-harming trials, participants exhibited increased activation in the orbitofrontal cortex (OFC), anterior mid-cingulate cortex (aMCC), amygdala, anterior insula (aIN), ventromedial prefrontal cortex (vmPFC), posterior superior temporal sulcus/temporal parietal junction (pSTS/TPJ), dorsolateral prefrontal cortex (dlPFC), and dorsal medial prefrontal cortex (dmPFC) during harming trials (Fig. 3B). There was no increase in activity for the non-harming conditions. For tool scenarios, stronger activation was observed in the aIN and pSTS/TPJ compared to hand scenarios. Conversely, the aMCC showed more pronounced activation for hand scenarios. Interaction effects indicated increased activation in the aMCC, amygdala, and dlPFC (Table 1).

**Fig. 3:**
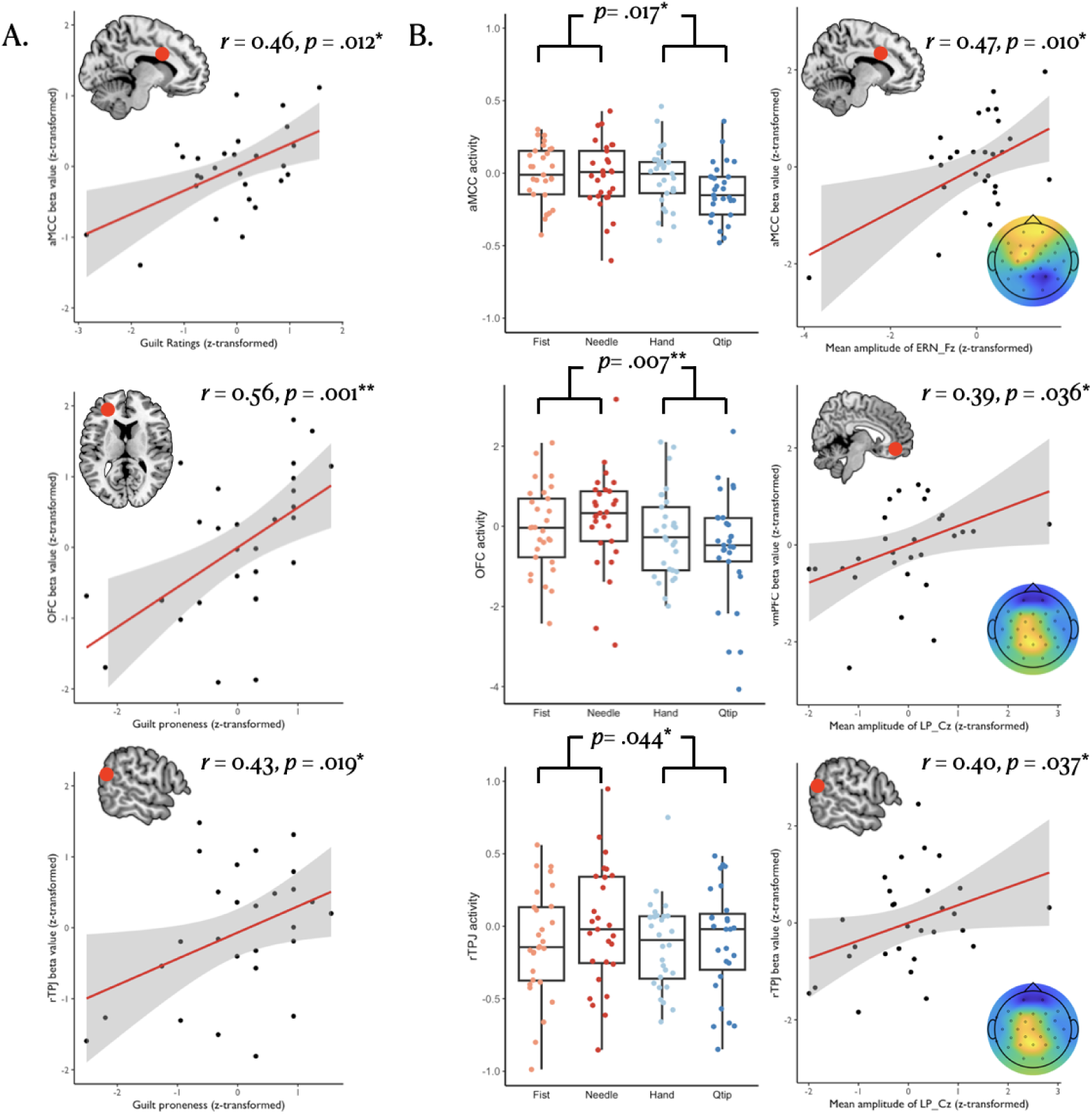
fMRI Correlations with Behavioral and ERP Results. **A.** Subjects’ self-reported guilt ratings positively correlate with aMCC activity (r = 0.46, p = 0.012). Additionally, their dispositional guilt trait (guilt-proneness) is positively associated with activity in the OFC (r = 0.56, p = 0.001) and right TPJ (rTPJ) (r = 0.43, p = 0.019). Both brain activities and behavioral ratings are normalized to z-scores for analysis. **B.** Bar plots depict variations in brain activity across different conditions. Notably, participants exhibited increased activity in the aMCC (p = 0.017) during harming trials, which positively correlated with their ERN amplitude (r = 0.47, p = 0.010). The OFC (p = 0.007) and rTPJ (p = 0.044) also demonstrated heightened activity in harming trials, alongside significant correlations with LP amplitude (OFC: r = 0.39, p = 0.036; rTPJ: r = 0.40, p = 0.037).

**Table 1:** Diverse BOLD responses activated under different moral conditions - ROI analysis results. Clusters marked with an asterisk are with a family-wise error (FWE) rate set at p < .05.

| Brain Region | Side | X | Y | Z | t |
| --- | --- | --- | --- | --- | --- |
| **Harm > No Harm** |  |  |  |  |  |
| OFC | L | -26 | 44 | 18 | 1.78 |
| OFC | R | 16 | 48 | 8 | 2.01 |
| aMCC | L | -14 | 8 | 30 | 1.84 |
| aMCC | R | 16 | -2 | 44 | 2.23 |
| Amygdala | L | -22 | -2 | -14 | 1.71 |
| aIN | R | 32 | 16 | 12 | 2.01 |
| vmPFC | L | 12 | 40 | -14 | 1.69 |
| pSTS/TPJ | R | 50 | -50 | 18 | 1.76 |
| dIPFC | R | 52 | 26 | 14 | 1.77 |
| dmPFC | R | 8 | 48 | 42 | 1.67 |
| **No Harm > Harm** |  |  |  |  |  |
| n.s. |  |  |  |  |  |
| **Tool > Hand** |  |  |  |  |  |
| aIN | L | -36 | 18 | 8 | 1.79 |
| pSTS/TPJ | R | 50 | -50 | 14 | 1.72 |
| **Hand > Tool** |  |  |  |  |  |
| aMCC | L | -16 | 10 | 32 | 1.73 |
| **(Needle + Touch) > (Q tip + Fist)** |  |  |  |  |  |
| aMCC | R | 10 | 2 | 30 | 1.67 |
| Amygdala | L | -22 | -4 | -20 | 2.10 |
| dIPFC | L | -48 | 34 | 28 | 1.71 |
| **(Needle + Touch) > (Q tip + Fist)** |  |  |  |  |  |
| aMCC | L | -16 | 6 | 44 | 1.98 |
| aMCC | R | 18 | 3 | 38 | 1.87* |

Abbreviations: R, Right; L, left; OFC, orbitofrontal cortex; aMCC, anterior midcingulate cortex; aIN, anterior insula; vmPFC, ventromedial prefrontal cortex; pSTS/TPJ, posterior superior temporal sulcus/temporal parietal junction; dIPFC, dorsolateral prefrontal cortex; dmPFC, dorsomedial prefrontal cortex.

To determine the impact of guilt-proneness and subjective guilt ratings on participants’ neural responses in harming conditions, an ROI analysis was performed focusing on specific guilt-associated brain regions. A participant’s dispositional guilt positively predicted activity in the OFC (r = 0.56, p = .001) and pSTS/TPJ (r = 0.43, p = .019). Meanwhile, activity in the aMCC (r = 0.46, p = .012) was the only region to significantly correlate with individuals’ subjective guilt ratings (Fig. 3A) (Table 2).

**Table 2:** Behavioral ratings correlate with unique BOLD clusters in the harming scenario.

| Brain Region | Side | MNI Coordinates |  |  | Peak t |
| --- | --- | --- | --- | --- | --- |
|  |  | X | Y | Z |  |
| Guilt proneness |  |  |  |  |  |
| OFC | L | -28 | 42 | 12 | 3.55 |
| pSTS/TPJ | R | 60 | -56 | 20 | 2.51 |
| Guilt ratings |  |  |  |  |  |
| aMCC | L | -12 | 8 | 44 | 2.69 |

Considering that subjects underwent two separate functional neuroimaging sessions, we performed regression analyses between mean ERP amplitudes and individual mean parameter estimates within pre-selected ROI regions to explore their alignment in immoral (harming) contexts. We found that activity in the anterior mid-cingulate cortex (aMCC) significantly correlated with ERN amplitude at both Fz (r = 0.47, p = .010) and Cz (r = 0.45, p = .014) (Fig. 3B). Similar patterns emerged for the anterior insula, which showed significant correlations with ERN amplitude at Fz (r = 0.44, p = .018) and Cz (r = 0.45, p = .037). Additionally, the mean amplitude of LP at Fz was significantly associated with amygdala activity (r = 0.38, p = .040), and the right temporal-parietal junction (TPJ) demonstrated a positive correlation with LP amplitude, albeit with marginal significance (r = 0.36, p = .053). At Cz, significant correlations were observed between LP amplitude and activity in the amygdala (r = 0.41, p = .026), ventromedial prefrontal cortex (vmPFC) (r = 0.39, p = .036), and right TPJ (r = 0.40, p = .037) (Fig. 3B) (Table 3).

**Table 3:** The mean amplitude of event-related potentials in harmful scenarios correlates with specific BOLD responses.

| Brain Region | Side | MNI Coordinates |  |  | Peak t |
| --- | --- | --- | --- | --- | --- |
|  |  | X | Y | Z |  |
| ERN at Fz |  |  |  |  |  |
| aMCC | L | -20 | 8 | 30 | 2.75 |
| aIN | L | -34 | 14 | 14 | 2.52 |
| ERN at Cz |  |  |  |  |  |
| aMCC | L | -20 | 8 | 30 | 2.64 |
| aIN | L | -36 | 10 | 14 | 2.10 |
| LP at Fz |  |  |  |  |  |
| Amygdala | L | -28 | -6 | -22 | 2.15 |
| pSTS/TPJ | R | 53 | -54 | 10 | 2.02 |
| LP at Cz |  |  |  |  |  |
| Amygdala | L | -18 | -8 | -14 | 2.36 |
| vmPFC | L | 6 | 42 | -14 | 2.2 |
| pSTS/TPJ | L | 52 | -52 | 12 | 2.05 |

## Discussion

The present study aimed to explore how individuals subjectively process the concept of wrongness and to what extent this is reflected by the neuropsychological mechanism of objective error awareness. Diverging from traditional psychological tasks that rely on absolute answers to gauge correctness, we employed a mental simulation task that encompasses moral and immoral situations, adapting the paradigm for EEG and fMRI studies, respectively. The neural activities recorded by both methods allow us to shed light on how individuals navigate the subjective perception of wrongness. Notably, compared to non-harming scenarios, a larger negative waveform was observed immediately following a harming action, indicating that participants respond more negatively to events they deem morally incorrect. Thus, the ERN does not merely reflect objective wrongness but also encompasses those actions we subjectively perceive as wrong.

By integrating EEG and fMRI results, we found that individuals’ ERN amplitude successfully predicted activity in their anterior midcingulate cortex (aMCC) and anterior insula (aIN). In our experiments, participants were asked to imagine themselves inflicting pain on others during harming trials. The aMCC is associated with negative affect, pain (Shackman et al., 2011; Tolomeo et al., 2016; Vogt, Berger, & Derbyshire, 2003), and experienced guilt (Cheng et al., 2021; McLatchie, Giner-Sorolla, & Derbyshire, 2016; Yu, Hu, Hu, & Zhou, 2014) . The anterior cingulate cortex was also involved in conflict monitoring during moral judgments (Greene, Nystrom, Engell, Darley, & Cohen, 2004). Moreover, several studies have shown that the ERN should originate from the anterior cingulate cortex (Herrmann et al., 2004; Markus Ullsperger, Danielmeier, et al., 2014). As for the anterior insula, it plays a crucial role in general aversive processing related to pain (Coghill, Sang, Maisog, & Iadarola, 1999; Starr et al., 2009) and empathy for pain (Singer et al., 2004). Furthermore, (Gu et al., 2012). Besides, M. Ullsperger, Harsay, Wessel, and Ridderinkhof (2010)) reviewed the role of the aIN in error processing and awareness, concluding that the aIN is essential for awareness of errors in automatic responses. In conditions involving financial harm, both the anterior cingulate cortex and the aIN showed increased activity (Greening et al., 2014). Additionally, Bastin and colleagues (2016), using stereo-EEG, found a consistent ERN response in both the aIN and ACC across all participants. Our findings resonate with previous research, highlighting the connection between ERN and specific brain regions, and further suggesting the automatic processing of error awareness and the early stages of moral judgment.

On the other hand, we discovered that participants’ guilt proneness was correlated with their ERN amplitude in harming scenarios, with individuals prone to guilt demonstrating larger ERN amplitudes. Given that guilt arises from wrongdoing and encourages avoidance of transgression (Baumeister, Stillwell, & Heatherton, 1994), individuals with higher guilt proneness were more inclined to steer clear of morally wrong behaviors (Cohen et al., 2012). Thus, our findings imply that the ERN not only reflects our capacity to detect errors in moral situations but also highlights individual differences in assessing moral wrongness.

In later stages, participants exhibited larger LP amplitudes in harming scenaios, which significantly correlated with activity in their amygdala, ventromedial prefrontal cortex (vmPFC), and right temporoparietal junction (TPJ). The amygdala and vmPFC are key to affective processing within moral cognition (Koenigs et al., 2007; Pascual, Gallardo-Pujol, & Rodrigues, 2013). Shenhav and Greene (2014) found that the amygdala is involved in emotional assessment, whereas the vmPFC integrates signals in moral judgments, with amygdala-vmPFC functional connectivity being particularly strong in emotion-based moral judgments. Additionally, the right TPJ is engaged in empathetic processing (Morelli, Rameson, & Lieberman, 2014; Shamay-Tsoory, 2011), linking TPJ thickness to cognitive empathy by Massey et al. (2017). Moreover, Young, Camprodon, Hauser, Pascual-Leone, and Saxe (2010) demonstrated that TMS applied to the right TPJ led participants to view harmful actions as less immoral, underscoring the importance of the right TPJ in moral judgment. Therefore, we propose that LP amplitude reflects the integration of cognitive and affective processing in moral judgment.

Despite participants reporting a stronger sense of guilt in harming scenarios, our study did not identify a significant correlation between LP amplitude and self-reported guilt. This observation aligns with findings from Sánchez-García and colleagues (2019), who noted a pronounced sense of guilt in self-wrong scenarios but failed to establish a correlation between self-reported guilt and LP amplitude. Sánchez-García suggested that LP might not fully encapsulate guilt, arguing it would be improbable for such a complex emotion to be solely represented by a single ERP component. Echoing this perspective, our results suggest that LP encompasses the broader affective processing in moral judgment, capturing moral emotions like guilt but not limited to them.

Nevertheless, our study faces limitations that warrant discussion. A primary concern is ecological validity; although we asked participants to immerse themselves in the action through animated simulations, this method of perspective-taking diverges from real-life experiences. Additionally, the EEG and fMRI sessions were conducted on separate days, challenging the direct integration of these data sets. By focusing on specific components and regions identified in prior research and averaging data across trials for each condition (Scrivener, 2021), we contend that our approach of amalgamating independently recorded data suffices to yield reliable outcomes.

Taken together, in contrast to conventional error studies, the ERN extends beyond merely signaling the detection of errors in moral judgments; it reflects individual variability when encountering moral transgressions. Furthermore, the LP plays a role in the affective dimension of moral processing during later stages. Our research offers insights into the individual differences in moral judgments and provides neural evidence in support of a dynamic moral cognition system. Reinforcing the dynamic moral framework, our findings highlight individual variability in moral judgments, underscored by distributed and overlapping neural networks encompassing various cognitive functions. This indicates that our subjective assessments of moral wrongness are supported by neural mechanisms akin to those involved in objective error awareness.

